# Low levels of metabolic auxotrophy among environmental *Pseudomonas* isolates

**DOI:** 10.64898/2026.02.13.705048

**Authors:** Simon Maréchal, Benjamin Heiniger, Shaohua Gu, Swagatika Dash, Christian H. Ahrens, Rolf Kümmerli

**Affiliations:** Department of Quantitative Biomedicine, University of Zürich, Switzerland; Method Development and Analytics, Agroscope, Zürich, Switzerland; PhD Program Systems Biology, Zurich Life Sciences Graduate School; College of Resources and Environmental Sciences, State Key Laboratory of Nutrient Use and Management (SKL-NUM), National Academy of Agriculture Green Development, China Agricultural University, Beijing 100193, China; Department of Ecology, School of Biology/Chemistry, University of Osnabrück, Osnabrück, Germany; Swiss Institute of Bioinformatics

**Keywords:** amino acids auxotrophy, natural bacterial communities, alternative biosynthetic pathways, soil and pond habitats, bioinformatic prediction, phylogenetic associations

## Abstract

Auxotrophy, the inability of bacteria to synthesize one or multiple essential metabolites (e.g. amino acids, vitamins, metabolites) is thought to be common among bacteria. However, studies often rely either on bioinformatic tools to predict auxotrophies from genome data or on experiments with low numbers of strains. Here, we combine experimental and bioinformatic approaches to assess amino acid auxotrophy levels among 315 co-isolated natural *Pseudomonas* strains from pond and soil habitats. Both approaches revealed that *Pseudomonas* isolates are predominantly prototrophs. We identified one single histidine auxotroph and five non-specific auxotrophs featuring complex growth phenotypes incompatible with single amino acid auxotrophies. While different bioinformatic pipelines vary in the extent to which auxotrophy is over- or underestimated, none of the pipelines could resolve the basis of non-specific auxotrophies. Our analysis further revealed the existence of multiple alternative biosynthesis pathways for methionine, proline, and phenylalanine, with significant enrichments of specific pathways among soil or pond strains. We conclude that combining experiments with bioinformatics is a powerful approach to assess the metabolic potential of environmental bacteria. Moreover, taxa like *Pseudomonas* can be predominantly prototrophic possibly owing to their generalist lifestyle, thus calling for nuanced ecological concepts predicting auxotrophy levels based on lifestyle and habitat.

## Introduction

Most environments are populated by complex bacterial communities consisting of many different species (Gibbons and Gilbert, 2015; Sunagawa *et al*., 2015). Given that space and nutrients within these communities are typically limited, there are ample opportunities for bacteria to interact in either negative or positive ways (Foster and Bell, 2012; Ghoul and Mitri, 2016; Kehe *et al*., 2021; Figueiredo *et al*., 2022). Negative interactions are ubiquitous and typically involve indirect competition for resources or direct interference competition through various mechanisms including the release of toxins or the deployment of contact-dependent inhibition systems (Ghoul and Mitri, 2016; Granato *et al*., 2019). Positive interactions also occur and often manifest in the release of beneficial products and metabolites that can be used by other community members (West *et al*., 2007; Kost *et al*., 2023). Such cooperative interactions can be based on the sharing of extracellular enzymes, siderophores and biofilm components (Nadell *et al*., 2009; Garcia-Garcera and Rocha, 2020; Kramer, Özkaya, *et al*., 2020; Pontrelli *et al*., 2022), or can involve metabolic interactions (Schink, 2002; Morris *et al*., 2012; D’Souza *et al*., 2018; Kehe *et al*., 2021; Giri *et al*., 2022; Pauli *et al*., 2023; Hesse and O’Brien, 2024).

The basic premise of metabolic interactions is that not all bacterial species in a community need to encode all the essential metabolic pathways. Instead, different bacteria can specialize on specific pathways or pathway steps and release metabolites into the environment, which can then be used as nutrient sources by metabolically complementary species in the community (Johnson *et al*., 2012). Such metabolic interactions can foster dependencies between species as community members may lose specific pathways altogether and thus shift from an autonomous (prototrophic) to a dependent (auxotrophic) lifestyle (Morris *et al*., 2012; Seif *et al*., 2020). Dependencies can either be linear, whereby species B relies on the release of a metabolite from species A, or circular whereby species A and B mutually depend on the release of specific metabolites from the other species (Harcombe *et al*., 2018). Metabolic interactions can promote evolutionary genome reduction and streamlining whereby species lose one or several biosynthetic pathways and thereby become fully dependent on the use of external sources of the corresponding metabolites (Wolf and Koonin, 2013).

Studying metabolic interactions and the evolution of dependencies have become popular in recent years (D’Souza *et al*., 2014; Pande and Kost, 2017; Johnson *et al*., 2020; Kim *et al*., 2021; Starke *et al*., 2023; Kasalo *et al*., 2025). This is because metabolic interactions are regarded as key elements explaining microbial community diversity and stability, and in the context of hosts they have been associated with host health and disease (Ramsey *et al*., 2011; Goldford *et al*., 2018; Starke *et al*., 2023). However, current study approaches come with specific limitations. Large-scale metabolic interaction studies are typically based on bioinformatic modelling (Price *et al*., 2020; Seif *et al*., 2020; Zimmermann *et al*., 2021). Although these approaches are powerful to predict metabolic interactions in complex communities, experimental validation is often not possible. Conversely, empirical studies provide solid experimental data but are often restricted to a relatively low number of (culturable) species or strains (Harcombe *et al*., 2018; Giri *et al*., 2021). Consequently, we still have a limited understanding of how common auxotrophies and metabolic dependencies are among strains and species in natural communities, and how good the match between bioinformatic approaches and empirical data is.

In our work, we aim to address these two open questions by connecting empirical experiments with bioinformatic analysis. Specifically, we take advantage of a well-characterized collection of 315 *Pseudomonas* isolates sampled from two habitats (soil and pond) (Butaitė *et al*., 2018; Kramer, López Carrasco, *et al*., 2020) to examine the ability of the isolates to synthesize proteinogenic amino acids. First, we assessed the frequency and specificity of amino acid auxotrophy using culture-based approaches. Second, we used different bioinformatic tools to screen the genomes of all isolates to detect amino acid biosynthetic pathways, the overall diversity of pathways, and potential gaps therein. Third, we compared experimental with bioinformatic data to examine the level of consistency across the approaches. Finally, we tested whether the abundance of specific amino acid pathways correlates with habitat type (soil versus pond) or phylogeny of the isolates.

## Results

### Natural *Pseudomonas* isolates show low levels of amino acid auxotrophy

In a first assay, we grew each of the 315 strains in either M9 minimal medium with glucose (M9G) but without amino acids or in M9G supplemented with casein hydrolysate (a source of proteinogenic amino acids) (Fig. 1A). We expect prototrophs to grow well in both media, whereas auxotrophs should only be able to grow in the presence of supplemented amino acids (Fig. 1B). When qualitatively comparing the growth across the two media, we found that 92.4 % (291 of 315) of all isolates grew well in the absence of supplemented amino acids, suggesting that these isolates are amino acid prototrophs (Fig. 1C). Among the remaining 24 isolates, only 5 isolates displayed a complete absence of growth in the amino acid depleted medium whereas the remaining 19 isolates showed reduced growth compared to the medium with amino acids (Fig. 1C). These results suggest that only a minority of natural *Pseudomonas* isolates are amino acid auxotrophs.

**Figure 1.**
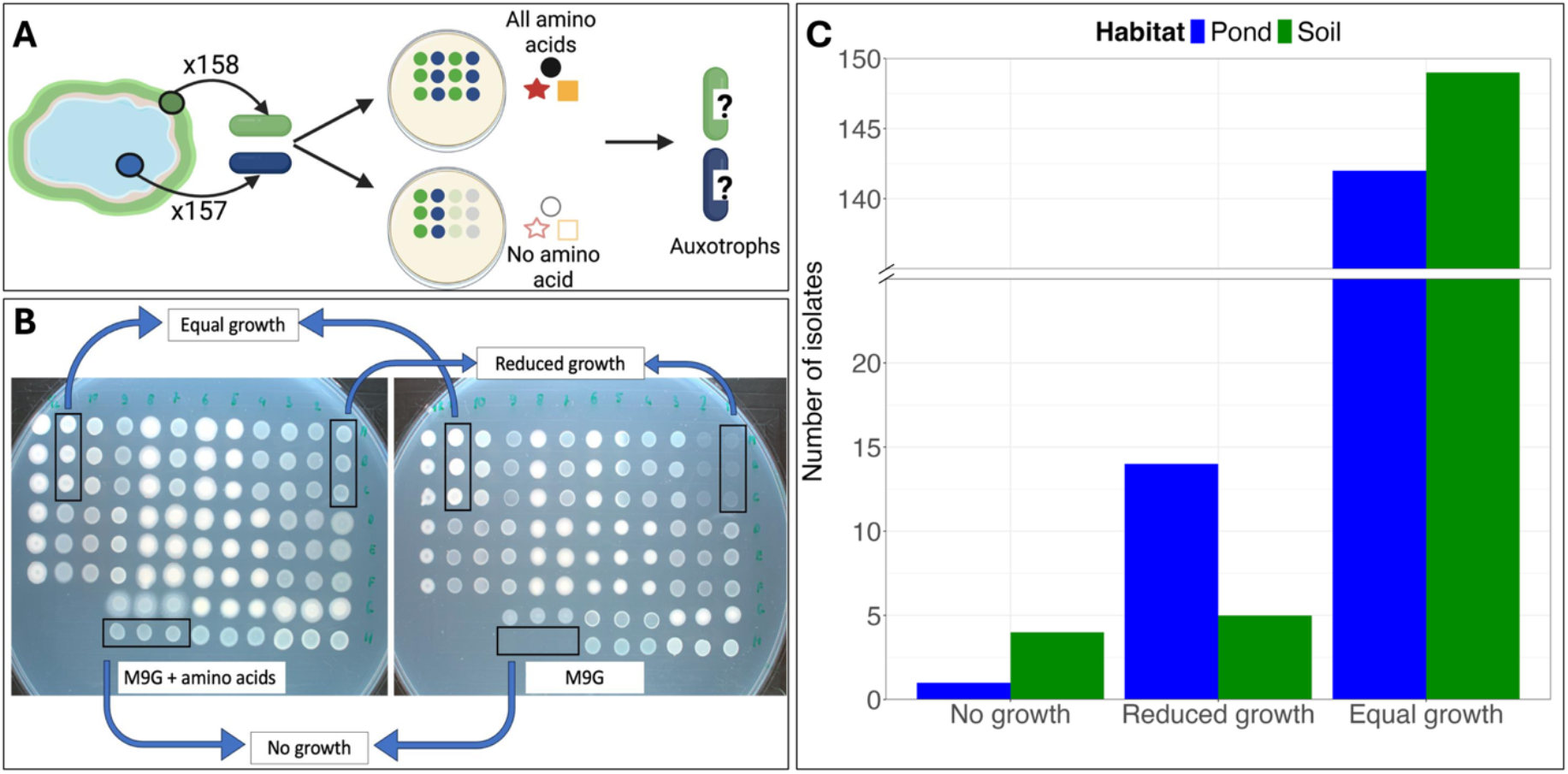
Amino acid growth assay revealed low levels of auxotrophy among natural *Pseudomonas* isolates: **A)** 315 natural *Pseudomonas* isolates, originally sampled from pond and soil habitats, were grown on M9G containing either all or none of the proteinogenic amino acids to detect putative amino acid auxotrophies. **B)** Representative experimental plates showing the growth of the same set of environmental isolates on the two different M9G media in triplicates. Growth between media was compared qualitatively and categorized as follow: “equal growth” when no growth difference was observed between the two media, “reduced growth” when the growth was reduced on the medium without amino acids, and “no growth” when no detectable growth could be observed on the medium without amino acids. Growth phenotypes were highly consistent across the three replicates and time (24h vs. 48h). **C)** Number of isolates in each growth category split according to habitats (blue = pond, green = soil).

### Single amino acid addition or omission experiments identified one specific auxotrophic isolate

Next, we focused on the 24 isolates displaying reduced or no growth in the absence of amino acids in the medium to explore their potential auxotrophy in more detail. As before, we used M9G medium, but this time we supplemented it with either one single amino acid (single addition, SA) or omitting a single amino acid from the mix of 20 (single omission, SO). This experimental design allows us to distinctively identify specific amino acid auxotrophies. For instance, a tryptophan auxotroph would grow in M9G supplemented with tryptophan as the sole amino acid source but would not grow in M9G with 19 amino acids but lacking tryptophan.

To validate our approach, we used a subset of 12 *Escherichia coli* mutants from the KEIO collection (Mee *et al*., 2014) where each mutant was deficient for the synthesis of a single specific amino acid. When growing these mutants in single omission M9G media, we found that all 12 *E. coli* mutants failed to grow in the medium lacking the respective amino acid, as expected (Fig. S1, left panel). Curiously, all these mutants also failed to grow in M9G without isoleucine, a pattern that cannot simply be explained by the mutational background of these strains (see Fig. S1 for an in-detail treatment of this case). We further performed the single amino acid addition experiment with two *E. coli* mutants (Δ*metA* and Δ*ilvA*) and observed (as expected) that their growth was recovered when M9G was supplemented with methionine and isoleucine, respectively (Fig. S1, right panel). Overall, these results demonstrate that our approach works and show that single amino acid auxotrophies can reliably be identified with our SA/SO approach.

With this prerequisite established, we conducted the SA and SO experiments with the 24 *Pseudomonas* isolates that showed reduced or no growth in our initial screen. We found that 18 isolates showed consistent growth across all media compositions, suggesting that they are in fact prototrophs (Fig. 2). Among the remaining 6 isolates, we found strong evidence for a histidine auxotrophy in isolate s3h05. Conversely, the other five isolates (s3a20, s3g13, s3h13, s3g15, p3F08) grew all perfectly fine in every SO medium but were either unable to grow (s3a20) or showed variable binary growth patterns across SA media (Fig. 2). We concluded that these isolates are non-specific auxotrophs. Overall, the amino acid SA and SO experiments strengthened the interpretation that most *Pseudomonas* isolates are prototrophs. Only 6 isolates (1.9%) showed growth patterns compatible with auxotrophy.

**Figure 2.**
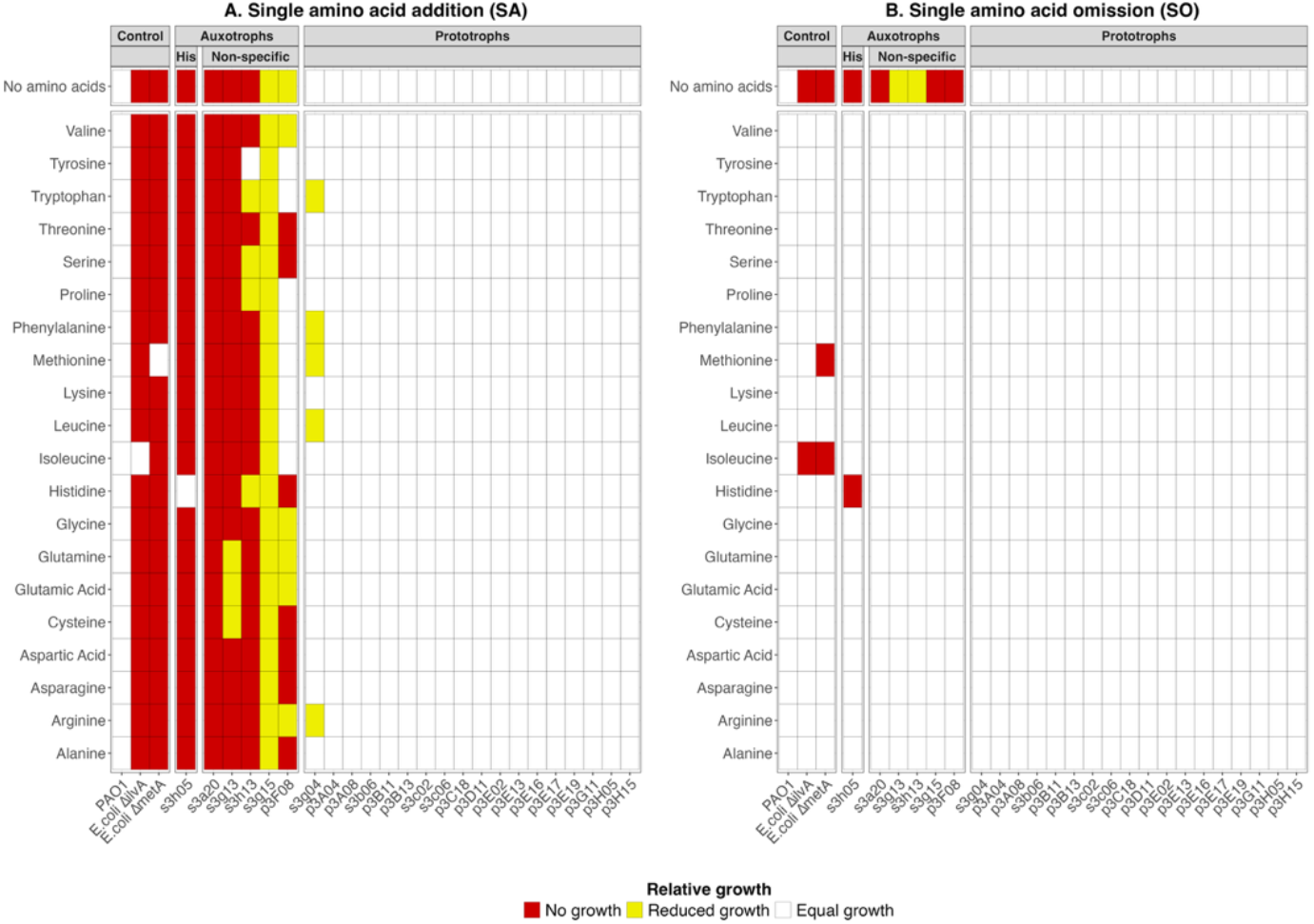
Growth assays on SA and SO medium confirmed a single specific and five non-specific auxotrophs: The 24 *Pseudomonas* isolates that showed no or reduced growth in the initial screen (Fig. 1) were subjected to two specific assays with the aim to identify the specificity of their putative amino acid auxotrophy. Relative growth values were assessed by comparing the growth of each isolate (x-axis) in medium containing all amino acids to media to which either a single amino acid was added (**A**, SA condition) or from which a single amino acid was omitted (**B**, SO condition). The strains are grouped from left to right into three categories. 1. Control strains: consisting of the prototroph *P. aeruginosa* PAO1 and the auxotrophs *E*.*coli*ΔilvA and *E. coli*ΔmetA. 2. Natural *Pseudomonas* isolates displaying auxotrophic phenotypes for either a specific single amino acid or an undefined set of amino acids (non-specific auxotrophs). 3. Natural *Pseudomon*as isolates with reduced growth in the initial screen (Fig. 1) but displaying prototrophic growth phenotypes in the SA/SO experiments. The focal amino acids, added to or omitted from M9G, are shown on the y-axis.

### Bioinformatic analysis confirm that most *Pseudomonas* isolates are prototrophs

To complement our experimental data, we conducted bioinformatic analyses on the sequenced genomes of 314 *Pseudomonas* natural isolates (one isolate was excluded from the analysis due to poor sequencing quality). We aimed to (i) create an inventory of all amino acid synthesis pathways present in each isolate, (ii) identify incomplete pathways that could explain the observed auxotrophies, and (iii) check for the presence of alternative pathways for the same amino acid across isolates. For these analyses, we used GapMind (Price *et al*., 2020), a software that can predict biosynthetic pathways of 17 amino acids and putative gaps therein with a confidence score (Fig. 3, Fig. S2). Notably, GapMind cannot predict biosynthetic pathways for alanine, aspartate and glutamate. This is because these pathways involve intermediates from central metabolism and non-specific transaminases that are generally present in genomes. Thus, it is generally assumed that bacteria can produce these three amino acids (Price *et al*., 2020). As prediction confidence can vary depending on the annotation method used, we ran GapMind on our genomes annotated with three different pipelines: Prokka, NCBI Prokaryotic Genome Annotation Pipeline (PGAP) and NCBI ORFfinder (ORFfind).

**Figure 3.**
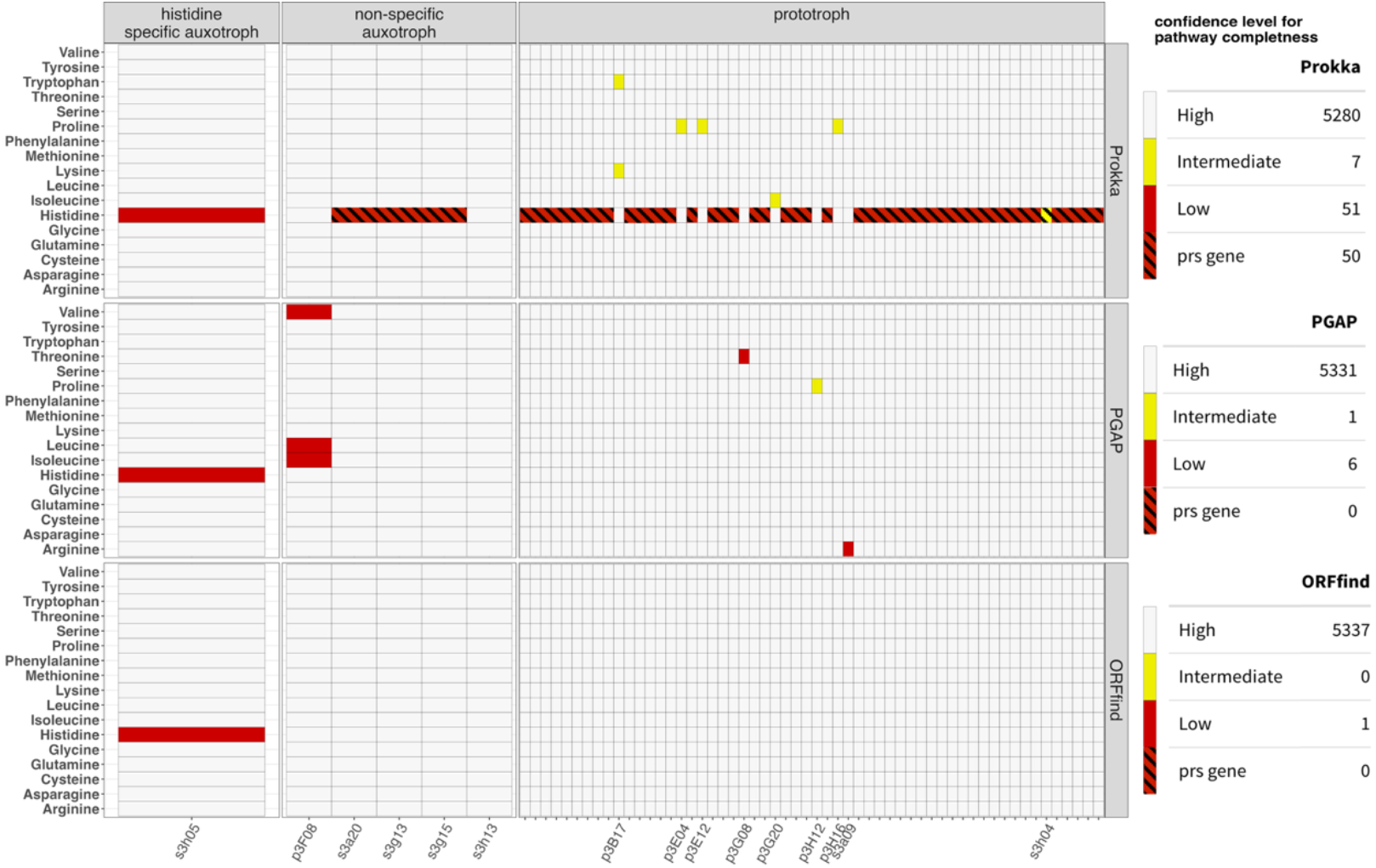
Comparing three bioinformatic pipelines to predict gaps in amino acid synthesis pathways: GapMind was used to predict amino acid synthesis pathways and possible gaps therein, from whole genome sequence data of 314 natural *Pseudomonas* isolates. The plot shows predicted gaps based on three different annotation pipelines (top row = Prokka, middle row = PGAP, bottom row = ORFfind). Isolates are ordered based on their experimentally determined amino acid metabolic phenotypes (from left to right: prototrophs, histidine-specific auxotroph, non-specific auxotrophs). Isolates with complete predicted pathways for all amino acids are excluded from this figure but can be found in Fig. S2. For prototrophs, isolate names are only shown for those with a predicted gap other than the *prs* gene (red-black hatched boxes). Tables on the right show the numbers of predicted amino acid synthesis pathways across all 314 isolates, stratified by annotation pipeline and prediction confidence: high (light gray), intermediate (yellow) and low (red). Low confidence predictions for *prs* (red-hatched) are shown separately. A critical point is that the confidence levels refer to the presence of a gene, such that low confidence for a pathway stands for high confidence for a gap.

All three pipelines confirmed that most *Pseudomonas* isolates are prototrophs. Specifically, we detected 5280, 5331 and 5337 complete amino acid synthesis pathways with high confidence among the 5338 total expected hits (314 isolates x 17 proteinogenic pathways) with Prokka, PGAP and ORFfind, respectively (Fig. 3). Accordingly, putative gaps were rare. Prokka predicted 58 gaps, whereas PGAP and ORFfind predicted only 7 and 1 gaps, respectively (Fig. 3). Only one gap was consistently predicted across the three annotation pipelines. It concerned a single gene (*hisF*), which encodes an imidazole glycerol phosphate synthase involved in histidine synthesis (Fig. S2). The isolate s3h05 that was predicted to lack *hisF* was indeed the specific histidine auxotroph identified in our experiments, thus yielding a perfect match between experimental and bioinformatic data.

Annotation with Prokka returned 50 putative gaps in the gene *prs* encoding a ribose phosphate pyrophosphokinase involved in the histidine biosynthesis pathway (Fig. S2). Since these gaps were not predicted by ORFfind and PGAP, we suspected them to be false positives. Indeed, all isolates with a predicted *prs* gap were prototrophs in our experiments. Moreover, the inflation in *prs* gap predictions with Prokka is a known problem for *Pseudomonas fluorescens* strains (Price *et al*., 2020).

With the PGAP annotation as basis, GapMind predicted five candidate gaps (in addition to *hisF*). One gap each was predicted for the arginine (isolate s3a09) and the threonine (isolate p3G08) pathways. We considered them as false positives because these predictions were only supported by one annotation method and moreover concerned isolates that had a prototrophic phenotype. The three remaining gaps all concerned the isolate p3F08 (a non-specific auxotroph) and a single gene (*ilvC*) encoding an enzyme connecting the synthesis pathways of isoleucine, leucine and valine. If this is a true positive the p3F08 isolate should be unable to grow in media that simultaneously lack isoleucine, leucine and valine. An experiment in M9G medium lacking these three amino acids, however, showed that p3F08 was not impaired in its growth, again suggesting that the predictions are false positives (Fig. S3).

In summary, our bioinformatic analyses reinforced the observed high levels of prototrophy among environmental *Pseudomonas* isolates and confirmed the only experimentally identified single amino acid auxotroph. In addition, we found that bioinformatic pipelines differ in their extent to which they generate false positive predictions and that they were unable to unravel the mechanistic basis of non-specific auxotrophies.

### GapMind reveals alternative pathways for methionine, proline and phenylalanine biosynthesis

We further used GapMind to detect alternative amino acid biosynthesis pathways encoded among our *Pseudomonas* isolates. The rationale for this analysis is to capture the metabolic diversity among pseudomonads. We found alternative pathways for methionine, proline, and phenylalanine biosynthesis (Fig. 4, Fig. S2).

**Figure 4.**
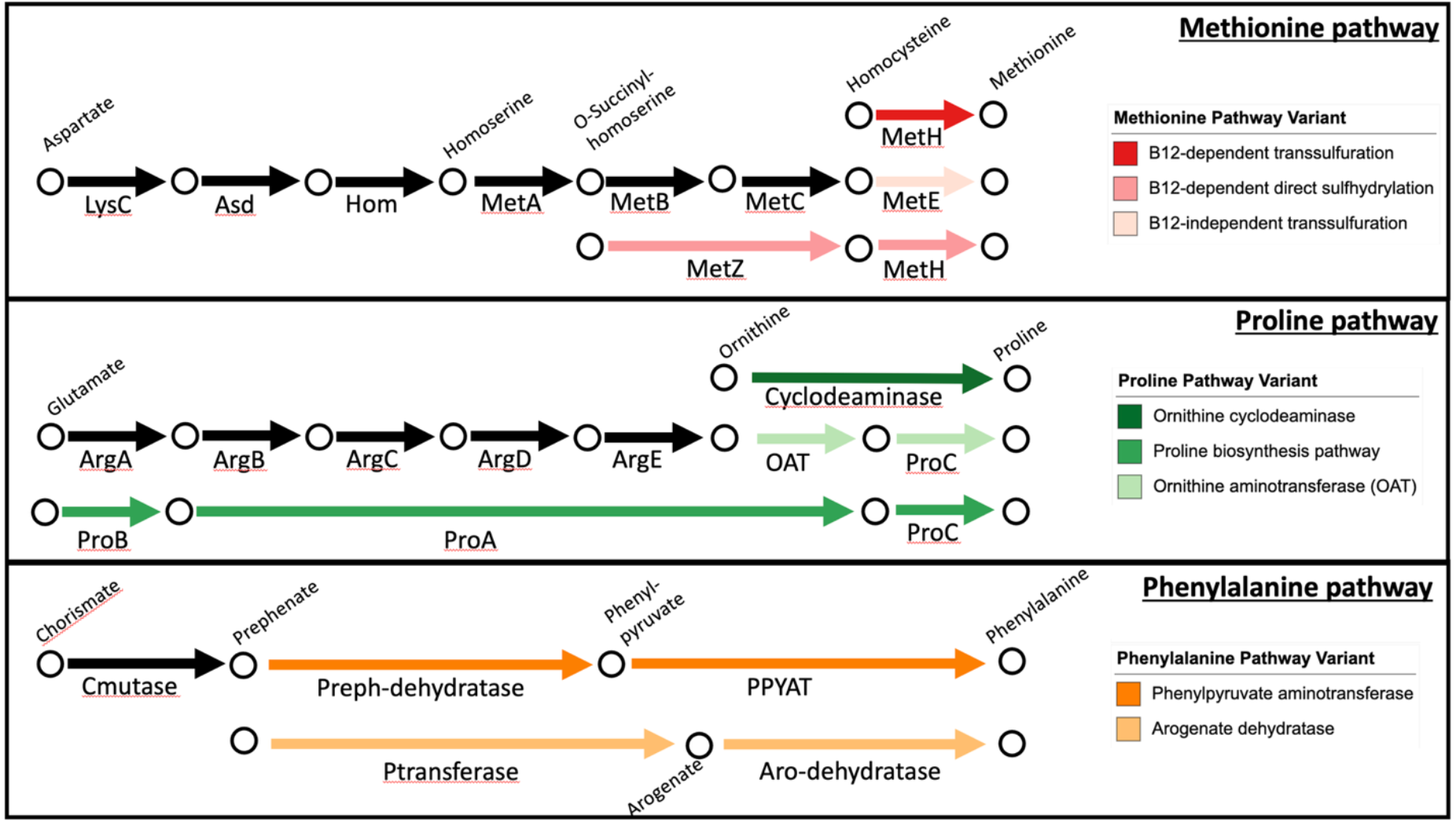
Alternative metabolic pathways for methionine, proline and phenylalanine. Shown are the alternative pathways for methionine, proline, phenylalanine biosynthesis encoded among the 314 natural *Pseudomonas* isolates. While two strains can differ in the pathway they possess, none of the strains featured two alternative pathways for the same amin acid. Black arrows represent steps that are shared among at least two pathways, whereas colored arrows show steps unique to each pathway. Biosynthetic pathways were reconstructed using KEGG and BioCyc database.

For methionine, the first four biosynthetic steps are conserved before the pathway splits into two branches, whereby O-succinyl-homoserine is converted to homocysteine either via the transsulfuration pathway (sulfur source: cysteine, enzymes: MetB and MetC) or the direct sulfhydrylation pathway (sulfur source: H_2_S; enzymes: MetZ). A second branching occurs in the transsulfuration pathway, in which homocysteine can be converted to methionine either via a vitamin B12-dependent (enzyme: MetH) or B12-independent (enzyme: MetE) pathway. We found that the B12-dependent transsulfuration was the most common pathway among the natural *Pseudomonas* isolates with a total of 278 occurrences (pond: 125, soil: 153, Table 1). The B12-dependent sulfhydrylation was the second most common pathway with 35 occurrences (pond: 31, soil: 4). Finally, we found only one occurrence of the B12-independent transsulfuration.

**Table 1.** Distributions of alternative amino acid biosynthetic pathways across soil and pond habitats and their association with phylogeny.

| Amino acid | Pathway variant | Number of variant |  | Association between pathways and habitats |  | Association between pathways and phylogeny |  |
| --- | --- | --- | --- | --- | --- | --- | --- |
|  |  | In pond | In soil | P-values of overall pathway distribution<sup>1</sup> | P-values of post-hoc pairwise comparisons<sup>2</sup> | Fritz & Purvis' D statistic | P-value for random phylogenetic structure<sup>3</sup> |
| Methionine | B12-dependent transsulfuration | 278 |  | < 0.0001 | < 0.0001 | 0.0214 | < 0.0001 |
|  |  | 125 (45%) | 153 (55%) |  |  |  |  |
|  | B12-dependent direct sulphydrylation | 35 |  |  | < 0.0001 | -0.0067 | < 0.0001 |
|  |  | 31 (88.6%) | 4 (11.4%) |  |  |  |  |
|  | B12 independent transsulfuration | 1 |  |  | / | / | / |
|  |  | 0 (0%) | 1 (100%) |  |  |  |  |
| Proline | Ornithine cyclodeaminsae | 169 |  | 0.0065 | 0.4230 | 1.006 | 0.5870 |
|  |  | 77 (45.6%) | 92 (54.4%) |  |  |  |  |
|  | Proline biosynthesis pathway | 137 |  |  | 1.0000 | 1.024 | 0.6903 |
|  |  | 71 (51.8%) | 66 (48.9%) |  |  |  |  |
|  | Ornithine aminotransferase | 8 |  |  | 0.0102 | 0.699 | 0.0129 |
|  |  | 8 (100%) | 0 (0%) |  |  |  |  |
| Phenylalanine | Phenylpyruvate aminotransferase | 238 |  | < 0.0001 | / | 0.187 | < 0.0001 |
|  |  | 103 (43.3%) | 135 (56.7%) |  |  |  |  |
|  | Arogenate dehydratase | 76 |  |  | / | 0.187 | < 0.0001 |
|  |  | 53 (69.7%) | 23 (30.3%) |  |  |  |  |

<sup>1</sup>Statistics based on Fisher's exact test on 2x3 contingency table with 10'000 Monte Carlo simulations
<sup>2</sup>Statistics based on Fisher's exact test on 2x2 contingency table, Bonferroni corrected for multiple testing
<sup>3</sup>Statistics based on Fritz & Purvis' D: null hypothesis (absence of phylogenetic signal): D = 1; alternative hypothesis (phylogenetic signal): D = 0.

For proline, there were two completely distinct pathways. The first biosynthetic pathway represents the standard proline synthesis route consisting of three enzymes (ProA-C). It was present in 137 isolates (pond: 71, soil: 66). The second biosynthetic pathway builds on the arginine pathway (enzymes: ArgA-E), which first yields ornithine. Ornithine is then converted to proline through either a cyclodeaminase or a two-step reaction involving the enzymes ornithine aminotransferase (OAT) and ProC (Fig. 4). The pathway involving the cyclodeaminase was more common being present in 169 isolates (pond: 77, soil: 92, Table 1), whereas the pathway variant involving the OAT was present in only 8 isolates (all pond).

For phenylalanine, there were two distinct pathways with a common first step involving a chorismate mutase generating prephenate. Subsequently, the pathway splits into two. In the first branch, prephenate is transformed into phenylpyruvate and finally to phenylalanine (enzymes: prephenate dehydratase, phenylpyruvate amino transferase (PPYAT)). In the second branch, prephenate is transformed into arogenate and then to phenylalanine (enzymes: prephenate amino transferase, arogenate dehydratase) (Fig. 4). The pathway involving phenylpyruvate as intermediate was more common occuring in 238 isolates (pond: 103, soil: 135, Table 1), whereas the pathway via arogenate occurred in 76 isolates (pond: 53, soil: 23).

### Alternative pathways are unevenly distributed across habitats and vary in the strength of their phylogenetic signal

We then tested whether the specific alternative pathways were enriched in soil versus pond habitats. Using Fisher’s exact test on contingency tables (Table 1), we found that alternative pathways were unevenly distributed between habitats (two-sided Fisher’s exact test, for methionine: p < 0.0001; for proline: p = 0.0065; for phenylalanine: p < 0.0001). For methionine, we found a significant enrichment of the B12-dependent transsulfuration pathway among soil isolates and a corresponding enrichment of the B12-dependent direct sulfhydrylation pathway among pond isolates (odds ratio, OR = 9.42, 95% CI: 3.2-37.7, Bonferroni-corrected p < 0.0001). For proline, the ornithine amino transferase pathway exclusively occurred among pond isolates (OR = 0, 95% CI: 0 - 0.6, Bonferroni-corrected p = 0.0102), whereas the other two pathways were more equally distributed among isolates of the two habitats (Table 1). For phenylalanine, there was an enrichment for the phenylpyruvate aminotransferase pathway among soil isolates and a corresponding enrichment of the arogenate dehydratase pathway among pond isolates (OR = 3, 95% CI: 1.7-5.5, p < 0.0001).

Next, we tested the strength of the phylogenetic signal for the alternative pathways. For this, we created a phylogenetic tree for our 314 natural isolates based on 1240 single copy orthogroups (Fig. 5, Fig. S4 for species prediction). We calculated the Fritz & Purvis’ D statistics for each of the alternative pathways using binary (presence vs. absence) data (Fritz and Purvis, 2010). The Fritz & Purvis D value captures the phylogenetic signal of each pathway on a sliding scale from random (D ≥ 1), to intermediate (0 < D < 1), to strong (D ≤ 0). Statistical tests indicate whether D-values are significantly different from 1 (non-random association: p_random_) (Table 1). For methionine, we found a strong phylogenetic signal for the B12-dependent transsulfuration (D = 0.021, p_random_ < 0.0001) and for the B12-dependent direct sulfhydrylation (D = ×0.007, p_random_ < 0.0001), meaning that alternative pathway distribution is almost entirely determined by phylogeny. For proline, we found the opposite pattern for all three pathway variants showing either random or very weak phylogenetic signals (ornithine cyclodeaminase pathway: D > 1, p_random_ = 0.5870; standard proline biosynthesis pathways: D > 1, p_random_ = 0.6903; ornithine aminotransferase: D = 0.699, p_random_ = 0.0129). For phenylalanine, both pathways showed a fairly strong phylogenetic signal (both phenylpyruvate pathway and arogenate pathway: D = 0.187, p_random_ < 0.0001).

**Figure 5.**
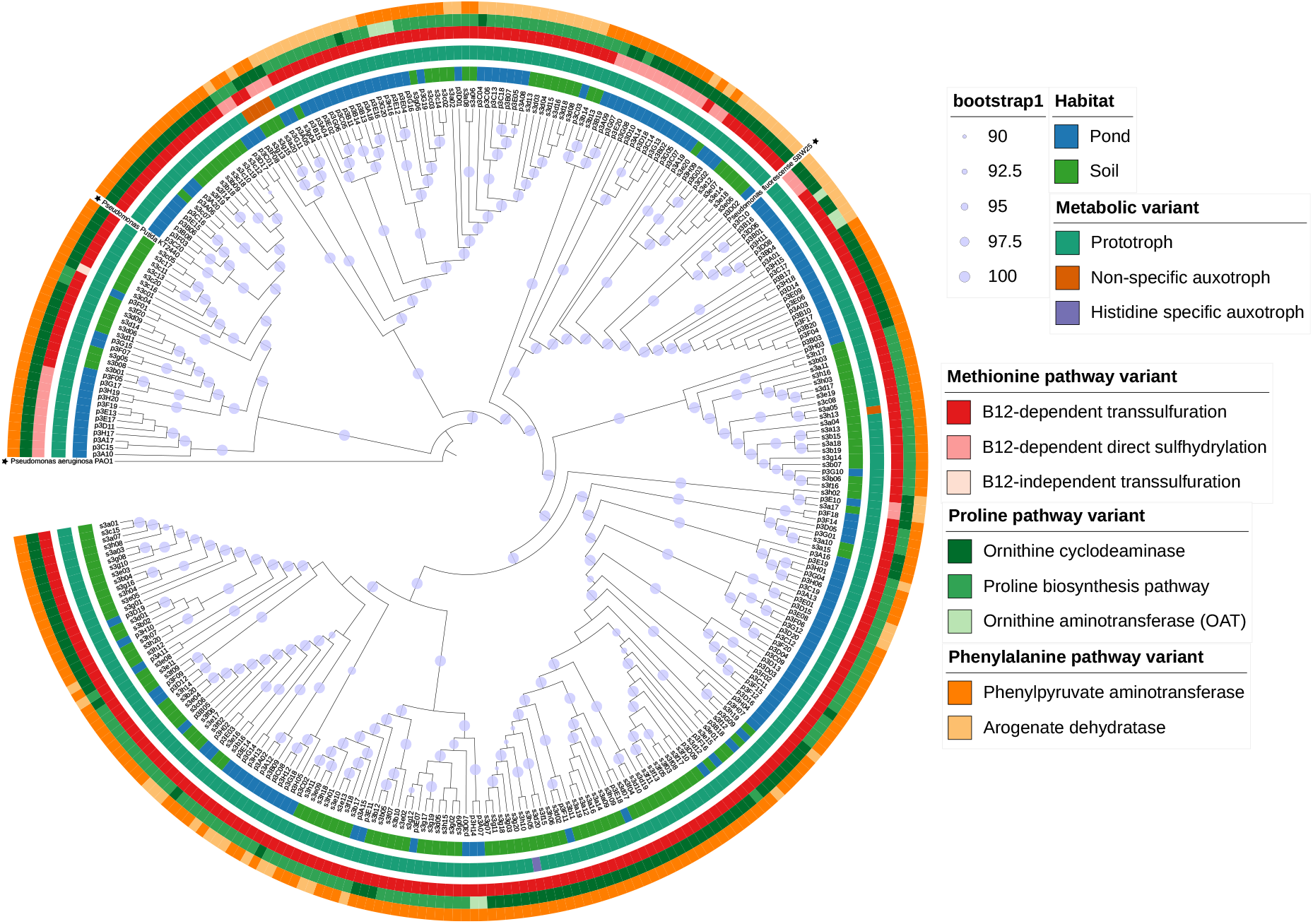
Strength of phylogenetic signal and habitat association vary across alternative amino acid biosynthetic pathways. The phylogeny of the 314 natural *Pseudomonas* isolates was assessed based on 1240 single copy orthogroups. *Pseudomonas aeruginosa* PAO1 was used as an outgroup, whereas *Pseudomonas putida* KT2440 and *Pseudomonas fluorescens* SBW25 were used as within-tree references (denoted by asterisks). Blue circles (in various sizes) show bootstrap values from 90% to 100%. From inside to outside, colored rings display the habitat of origin (soil vs. pond), the metabolic type (prototroph vs. auxotroph), and the alternative pathways for methionine, proline and phenylalanine. Alternative pathways show significant segregation according to habitat types. The phylogenetic signal is high for methionine and phenylalanine alternative pathways, but literally absent for proline alternative pathways (see Table 1 for statistics).

## Discussion

Metabolic auxotrophies, the inability to produce essential metabolites, are supposed to be common in microbial communities (Morris *et al*., 2012; Wolf and Koonin, 2013; D’Souza *et al*., 2014; Zengler and Zaramela, 2018; Johnson *et al*., 2020). The reason is that auxotrophy saves metabolic costs and thus generates fitness advantages as long as the essential metabolites can be acquired from the environment or from other community members overproducing and secreting them (D’Souza *et al*., 2018; Preussger *et al*., 2020; Kasalo *et al*., 2025). Auxotrophy can thus foster dependencies between microbial species and result in cross-feeding interactions. However, the frequency of auxotrophy is often inferred from sequencing data using global data bases (Price *et al*., 2020; Seif *et al*., 2020; Zimmermann *et al*., 2021; Ramoneda *et al*., 2023). Less clear is how common auxotrophies are among strains co-isolated from the same community and how well in silico predictions match empirical data. Here, we combined experimental and bioinformatic approaches to detect amino acid auxotrophies in a collection of 315 natural *Pseudomonas* strains, co-isolated from eight pond and eight soil communities. Our analyses revealed that experimental and bioinformatic approaches coincided in predicting high levels of prototrophy across habitats and communities. Specifically, both approaches identified only a single auxotrophic isolate for histidine. However, there were also several mismatches between the two approaches. While our experiments revealed five non-specific auxotrophs that were unable to grow in media without amino acids, in silico approaches were unable to identify the basis of these auxotrophies. Conversely, one of the three bioinformatic pipelines used systematically overestimated auxotrophies. Taken together, we found low levels of amino acids auxotrophies (1.9 %) among co-isolated *Pseudomonas* strains from pond and soil communities and show that a careful integration of experimental and bioinformatic approaches is required to reliably assess specific and non-specific auxotrophies.

At first sight, our findings contradict the perception that auxotrophies are ubiquitous in bacterial communities. However, a critical consideration is that the genus *Pseudomonas* is known for its metabolic and ecological versatility (Spiers *et al*., 2000; Silby *et al*., 2011). Given its ubiquity in many habitats and its generalist lifestyle, it is perhaps not surprising that the level of auxotrophy is low among *Pseudomonas* isolates (Ramoneda *et al*., 2023). In this context, we propose that the debate should not be about whether levels of auxotrophies are generally high or low, but more about gaining a better understanding of the ecological conditions that either favor high versus low levels of auxotrophies. The evolution of auxotrophy and thus metabolic dependencies requires stable interactions between cross-feeding partners. Stable interactions are more likely guaranteed when bacteria are adapted to specific niches and live in environments with little perturbations. Based on these considerations, we predict a negative correlation between auxotrophy frequency in a community and the proportion of taxa with a generalist (non-niche specific) lifestyle. Similarly, we expect a negative correlation between auxotrophy frequency in a community and the frequency of environmental perturbations.

Bioinformatic approaches are powerful to predict auxotrophies and metabolic interactions from (meta-) genome data and are especially useful to assess the metabolic potential of unculturable bacteria. However, our study shows that bioinformatic tools can both over and underestimate auxotrophies (see also Seif et al. 2020; Price et al. 2018). First, among the three different genome annotation tools used (Prokka, PGAP, ORFfinder), we found that Prokka tended to overestimate auxotrophy levels (Figure 3). The overestimation was predominantly associated with the non-annotation of a single gene (*prs*), which is part of the histidine synthesis pathway. Previous research reported an open reading frame for *prs* in *P. fluorescens* with 67% nucleotide sequence similarity with *E*.*coli* K-12 *prs* gene, but the annotation threshold used by Prokka is likely too stringent for the reliable annotation of this protein (Price *et al*., 2020). Second, we found that PGAP and ORFfinder tended to underestimate auxotrophies. Our experimental screens revealed five auxotrophic isolates that were unable to grow in media deprived of all amino acids. Our subsequent screens with SA and SO (single amino acid addition or omission) revealed that these isolates are non-specific auxotrophs. This means that these isolates cannot grow when a combination of several amino acids is missing but can synthesize each singly missing amino acid presumably through intertwined pathways. None of the bioinformatic pipelines could resolve the mechanistic basis of these non-specific auxotrophies. Although such non-specific auxotrophs seem to be common (D’Souza *et al*., 2014), our work shows that bioinformatic approaches fail to predict them and also experiments struggle to unravel details of this type of auxotrophy. Finally, previous bioinformatic work suggested that at least 40% of a synthesis pathway need to be missing for reliably classifying a strain as auxotroph (Ramoneda *et al*., 2023). This contrasts with our results that reliably identified (through experiments and bioinformatics) a histidine auxotroph based on a single gene deletion. Thus, such stringent bioinformatic thresholds could underestimate auxotrophies and cause mismatches between bioinformatic and experimental data. These considerations show that an integration of empirical and bioinformatic approaches is a powerful approach to assess the frequency and mechanistic basis of bacterial auxotrophies.

Our bioinformatic analysis further identified the presence of alternative biosynthesis pathways among isolates for methionine, proline, and phenylalanine. The distribution of these alternative pathways significantly correlated with the habitat type (pond vs. soil): strong association for methionine and phenylalanine; weaker association for proline. A strong habitat association could indicate that specific pathways are particularly beneficial in certain environments and may therefore be enriched in pond versus soil. In parallel, we observed strong phylogenetic signals for methionine and phenylalanine, yet almost no signal for proline. Strong phylogenetic signals indicate that certain amino acid biosynthetic pathways became established in specific *Pseudomonas* lineages and stayed there conserved over evolutionary timescales. Conversely, a weak phylogenetic signal indicates that a biosynthetic pathway can easily be lost or gained (through horizontal gene transfer) over evolutionary timescales. Importantly, we observed a positive collinearity between habitat effect and phylogenetic signal. It is therefore difficult to conclude whether the two habitats select for (i) different amino acid synthesis pathways or (ii) different species that tend to have different pathways. The only exception is the proline ornithine aminotransferase pathway, which only occurred among pond isolates with no significant phylogenetic signal (Table 1). Here, we can confidently conclude that this pathway is beneficial in aquatic environments regardless of the phylogenetic background of the strain. Altogether, we observed habitat-specific pathway enrichment, and it would be important to understand the ecological and evolutionary factors driving specificity not only in *Pseudomonas* but also in other taxa.

In conclusion, we combined experimental with bioinformatic approaches to show that most natural *Pseudomonas* isolates from pond and soil habitats are amino acid prototrophs. Although the match between bioinformatic predictions and experimental results was robust overall, there were certain bioinformatic pipelines that overestimated the frequency of auxotrophies, whereas others underestimated their occurrence. Mismatches predominantly involved non-specific auxotrophies, cases in which bacteria can only grow when multiple amino acids are supplemented. While previous studies reported auxotrophy to be common in natural communities, we here show that it can be rare in certain taxa, like *Pseudomonas*. Our results contribute to the development of general ecological principles explaining variation in levels of auxotrophies across taxa and habitats. In particular, we propose that environmental conditions (stability versus perturbation) and lifestyle (specialist versus generalist) are key characteristics determining whether levels of auxotrophies and metabolic dependencies are high or low, respectively.

## Material and methods

### Strain collection

We used a collection of 315 natural *Pseudomonas* strains isolated from pond and soil samples collected on Irchel campus of the University of Zurich (47.39° N, 8.54 °E), Switzerland (Butaitė *et al*., 2017). The initial sampling involved the collection of 8 different soil and 8 different pond samples, and the isolation of 20 *Pseudomonas* strains per sample. Among these 320 isolates, Sanger sequencing of the *rpoD* gene confirmed that 315 isolates belonged to the *Pseudomonas* genus and phylogenetic analyses show that most isolates cluster with either the *P. fluorescens* species complex, *P. putida* or *P. syringae* (Fig S5). Each isolate has an identification code, consisting of a habitat label (s = soil, p = pond), a location number (3 for Irchel campus), a community label (soil communities: lower case letters ‘a’ to ‘h’; pond communities: upper case letters ‘A’ to ‘H’) and an isolate number (1-20). We slightly modified the identification code compared to (Butaitė et al. 2018) by introducing the label ‘p’ for pond isolates.

As a prototrophic control strain, we used *Pseudomonas aeruginosa* PAO1. As auxotrophic control strains, we used a collection of 12 *Escherichia coli* mutants from the KEIO collection (Baba *et al*., 2006), each featuring a transposon insertion in a specific amino acid synthesis gene, rendering them auxotrophs for the corresponding amino acid (Table S1).

### Media preparation

For all experiments, we used M9 minimal medium supplemented with glucose (henceforth called M9G). Specifically, M9G consisted of pre-prepared 1x M9 salts mix (KH_2_PO_4_ (3 g/L), NaCl (0.25g/L), Na_2_HPO_4_ (6.78g/L), NH_4_Cl (1 g/L)), MgSO_4_ x 7H_2_O (2 mM), CaCl_2_ x 2H_2_O (0.1 mM), and glucose (0.4% m/v) as sole carbon sources. The M9 minimal medium was autoclaved while filter-sterilized (0.2μm) glucose and autoclaved MgSO_4_ x 7H_2_O and CaCl_2_ x 2H_2_O were added after autoclaving. We prepared five variants of M9G agar (1.5%) for the experimental amino acid auxotrophy screen: M9G agar (i) without amino acids, (ii) supplemented with an undefined mixture of all amino acids through the addition of 2g/L casamino acid (CAA, casein hydrolysate), (iii) supplemented with a defined mixture of all 20 amino acids, (iv) supplemented with a single amino acid (SA = single addition), and (v) supplemented with 19 out of the 20 proteinogenic amino acids (SO = single omission). Aqueous amino acid stock solutions were prepared at a final concentration of 20 mM, filter-sterilized (0.22μm) and stored at 4°C for not more than three months. Individual amino acids were filter-sterilized (0.2μm) individually and consequently mixed and added to M9G to a final concentration of 50 μM. All above ingredients were purchased from Merck (Buchs SG, Switzerland) except for the individual amino acid, which were purchased from neoFroxx GmbH (Germany).

### Preculture and culture conditions for the auxotrophy screen

We stored all isolates as glycerol stocks (25%) at -70°C. Before each screen, we revived the isolates by streaking them on Lysogeny broth (LB) agar plates, followed by incubation at 28°C for approximately 24h. From each plate, we picked a single colony to inoculate liquid precultures consisting of 1.5mL of M9G + CAA (see above), distributed in 24 well plates. Precultures were incubated at 28°C for 24h and 170 RPM. Subsequently, they were washed twice with a 1 x M9 salts solution followed by centrifugation at 9000 RPM for 2 min. The optical density (OD at 600 nm) of isolates was adjusted to 0.1. To prepare the experimental agar plates, we filled large round culture dishes (150 x 20 mm, Sarstedt, Germany) with 50mL of hot M9G agar. The version of the M9G agar used (i) - (v) dependent on the screen with the amino acid mixture being added after autoclaving as soon as the media was cooled down to approximately 60°C. For the initial auxotrophy screen with all 315 isolates, we used M9G agar versions (i) and (ii). For the in-depth screening of the putative auxotrophs, we used M9G agar version (i) and (iii) - (v).

For each screen, we spotted 2μL droplets of each OD-adjusted isolate using a Viaflo 96 automated pipetting system (Integra Biosciences, Switzerland). Each plate was seeded with triplicates of 28 environmental isolates, *P. aeruginosa* PAO1 (prototroph control) and *E. coli ΔilvA* (auxotroph control) and six medium blanks to check for contamination. Droplets were dried for 10 min under the hood and plates were then incubated at 28°C for 24 h. Subsequently, we imaged each agar plate using a FUSION FX6 EDGE Imaging System (Witec, Switzerland) and qualitatively assessed colony growth by comparing growth between media with and without (all or specific) amino acids. Specifically, colony growth on media containing all amino acids was taken as the standard for each isolate. Then, we compared this standard to colony growth of the corresponding isolate on media without (all or specific) amino acids. We defined three categories relative to the reference standard: equal growth, reduced growth and no growth. For all the isolates that showed reduced growth, we repeated the experiment and incubated for 48h to confirm that reduced growth is due to auxotrophy and not simply due to slow growth.

### Whole genome sequencing and GapMind analysis

We sequenced the whole genomes of all 315 isolated using Illumina short read sequencing technology. A subset of 24 isolates was sequenced as part of a separate study using Illumina Hiseq 2500 (2 × 250 bp paired end) and Shovill for genome assembly (a SPAdes based assembler)(Butaitė *et al*., 2017). The remaining isolates were sequenced in multiple batches by MicrobesNG (Birmingham, UK, https://microbesng.com/) using illumina NovaSeq 6000 (Illumina, San Diego, USA) (2 x 250 bp paired end, with 30x coverage). Here is a brief extract of the protocol used by Microbes NG. The genomic DNA libraries were prepared by MicrobesNG using a Nextera XT Library Prep Kit (Illumina, San Diego, USA) following manufacturer’s protocol with a modification to the DNA input (2-fold increase) and a PCR step (elongation time increased to 45 sec). Pipetting for DNA quantification and library preparation were done using a Hamilton Microlab STAR automated liquid handling system (Hamilton, Bonaduz AG, Switzerland). Adapter sequences were trimmed from reads using Trimmomatic (version 0.30) with a quality cutoff of Q15. Quality controlled was performed by MicrobesNG’s in-house pipelines (Samtools, BedTools, and bwa-mem). De novo genome assembly was also performed by MicrobesNG using SPAdes (version 3.7). Using these assembled genomes, we performed the genome annotation with three different tools: Prokka 1.14.3, NCBI PGAP 2024-04-27.build7426 (Li *et al*., 2021) specifying *Pseudomonas* sp. as the organism, and NCBI ORFfinder 0.4.3 (with genetic code 11 and default settings), which is not biased by homology-based gene prediction. We then used GapMind (Price *et al*., 2020) with PaperBLAST commit 93e68cf (Price and Arkin, 2017) to predict amino acid biosynthesis pathways and gaps therein, separately for each of the three annotations. We developed a Python script to summarize auxotrophies, whereas plots and data analysis were done with R (version = 4.4.3).

### Phylogenetic analysis

We assessed the orthology inference with OrthoFinder (Emms and Kelly, 2019) and further used the single copy orthogroups to reconstruct the phylogenetic association among the 314 sequenced *Pseudomonas* isolates. For orthology inference, we integrated three reference strains into our phylogenetic analysis: *P. aeruginosa* PAO1 (RefSeq: NC_002516.2) as an outgroup, and *P. fluorescens* SBW25 (RefSeq: NC_012660.1) and *P. putida* KT2440 (RefSeq: NZ_CP169744.1) as intra-tree references. We used all the identified single copy orthologue sequences (n=1240) to create the phylogenetic tree. In detail, these 1240 protein sequences were aligned using MAFFT (opt: --auto) and trimmed with trimAI (opt: -automated1). The trimmed aligned sequences were concatenated using a python script. Finally, IQ-TREE2 was used on the concatenated sequence with following parameter settings: -m LG+R7, -bb 1000, -alrt 1000, -nt 16. IQ-TREE2 ran on 317 sequences of 386854 aligned amino acids among which there were: 146841 distinct patterns, 145116 parsimony-informative, 26454 singleton sites, 215284 constant sites. To visualize the generated tree file, we used the tree visualization web platform iTOL (Letunic and Bork, 2024). To annotate the phylogenetic tree, we generated iTOL datasets using R.

### Statistical analysis

To assess whether pathways are differentially distributed across habitats, we first performed Fisher’s exact test on contingency tables [h x n] with “h” the number of habitats = “2” and “n” the number of alternative pathways per amino acid (n = 2 for phenylalanine and n = 3 for methionine and proline). For methionine and proline, where contingency tables were 2×3, we computed p-values with 10’000 Monte Carlo simulations as described in the R package {stats}. We then performed post-hoc Fisher’s exact tests for each of the alternative pathways (per amino acid) by comparing its frequency in pond and soil relative to the frequencies of the sum of the other pathways (2×2 contingency tables). We corrected p-values for multiple testing by using the Bonferroni approach.

We used Fritz and Purvis’ D statistics to quantify the phylogenetic signal of each alternative pathway. We considered alternative pathway abundance as a binary variable (present or absent). Statistical tests comprised 10,000 permutations and D-value < 1 with p_rand_ < 0.05 to reject the null hypothesis of random trait distribution. We used the Bonferroni approach to correct p-values for multiple testing. No statistical tests were performed for the alternative methionine pathway that occurred only once.

## Acknowledgments

We thank Christian Kost for comments on the manuscript. This project was funded by the Swiss National Science Foundation (Grant number 212266 to RK).

## Author contributions

Conceptualization: SM and RK; Experimental design: SM and SD; Experiments: SM; Data Analysis: SM, BH, SG; Writing (first draft): SM and RK; Writing (refinement): all coauthors

