## Supplementary data for "Low levels of metabolic auxotrophy among environmental *Pseudomonas* isolates"

**This file contains:**

- 1 Supplementary Table
- 4 Supplementary Figures

**Supplementary Tables**

**Table S1.** *E.coli* K12 mutants from the KEIO collection used in this study with a description of the mutated gene and its role in the respective amino acid biosynthesis pathway (from EcoCyc database). The mutants were previously described in Mee et al. 2019. Genome Biol 20: 238.

| **Gene name** | **Gene function (extracted from https://ecocyc.org/ECOLI/NEW-IMAGE?type=PATHWAY&object=Amino-Acid-Biosynthesis** |
|---|---|
| *argA* | N-acetylglutamate synthase *(*ArgA) carries out the first steps in the superpathway of arginine biosynthesis and the pathway of ornithine biosynthesis. |
| *cysE* | Serine acetyltransferase carries out the first step in the pathway of cysteine biosynthesis, converting L-serine into O-acetyl-L-serine. |
| *hisB* | HisB catalyzes two key reactions in histidine biosynthesis. |
| *ilvA* | Threonine deaminase (IlvA) carries out the first step in the synthesis of isoleucine. |
| *leuB* | catalyzes the first (LeuA)/third (LeuB) and second (LeuC) committed step in leucine biosynthesis |
| *lysA* | Diaminopimelate decarboxylase (LysA) catalyzes the last step in lysine biosynthesis |
| *metA* | (MetA) catalyzes the first unique step in the de novo methionine biosynthesis pathway in Escherichia coli |
| *pheA* | (PheA) carries out the shared first step in the parallel biosynthetic pathways for the aromatic amino acids tyrosine and phenylalanine, as well as the second step in phenylalanine biosynthesis. |
| *proA* | (ProA) catalyzes the the reduction of γ-glutamyl phosphate to glutamate 5-semialdehyde, the second reaction in the L-proline biosynthesis I (from L-glutamate) pathway |
| *thrC* | Threonine synthase (ThrC) carries out the final step in the biosynthesis of L-threonine |
| *trpC* | Bifunctional phosphoribosylanthranilate isomerase / indole-3-glycerol phosphate synthase (TrpC) carries out the third and fourth steps in the tryptophan biosynthesis pathway. |
| *tyrA* | Bifunctional chorismate mutase / prephenate dehydrogenase (TyrA) carries out the shared first step in the parallel biosynthetic pathways for the aromatic amino acids tyrosine and phenylalanine, as well as the second step in tyrosine biosynthesis. |

### Supplementary Figures

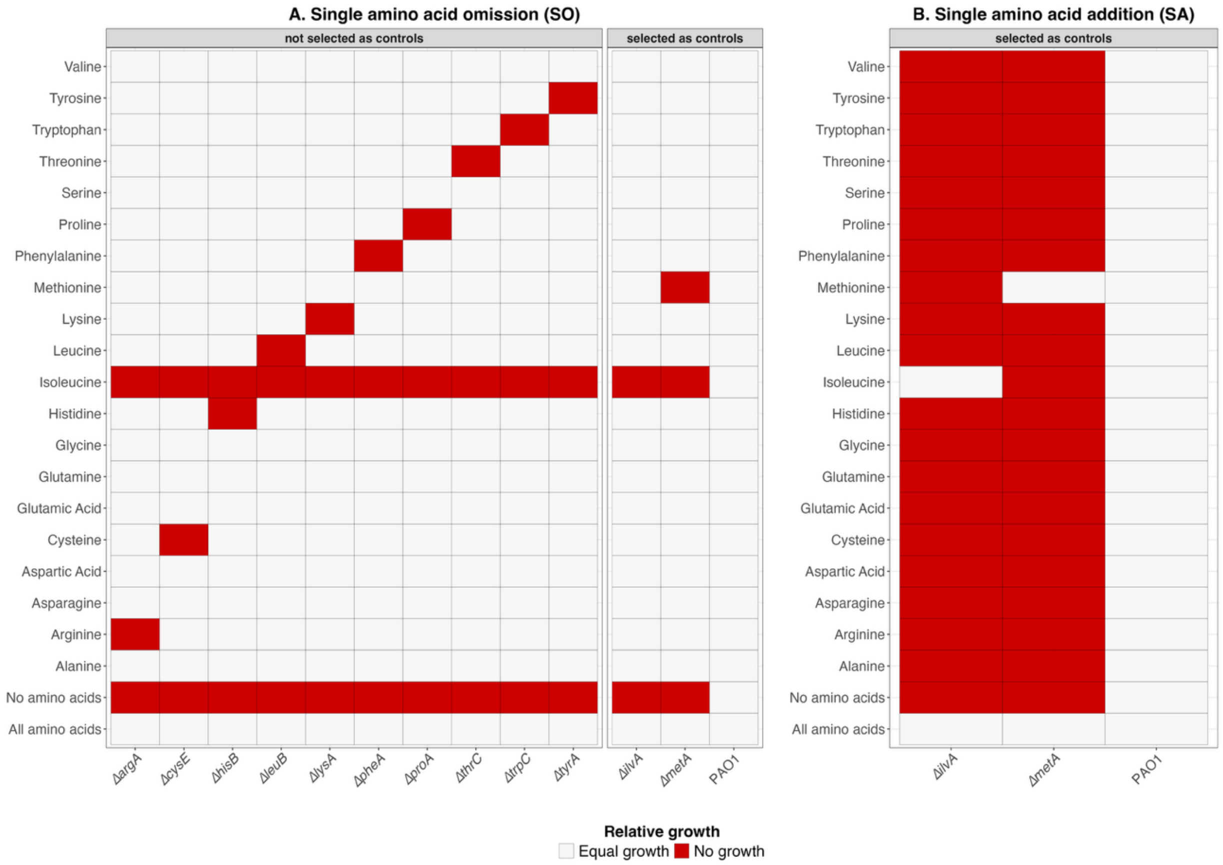


**Figure S1. Growth of the prototrophic *P. aeruginosa* PAO1 and the auxotrophic *E.coli* control strains used in this study**. **A)** The prototrophic PAO1 and 12 auxotrophic *E. coli* from the KEIO collection were used to validate our experimental approach. Each strain was grown in M9G agar with all amino acids and in M9G media supplemented with 19 out of the 20 proteinogenic amino acids (SO = single omission). See material and method for exact media composition. Data show growth of strains under SO conditions relative to M9G with all amino acids. **B)** The prototrophic PAO1 and two auxotrophic *E. coli* from the KEIO collection (*E.coli ΔilvA* and *E.coli* Δ*metA)* were grown in M9G with all amino acids and M9G supplemented with a single amino acid (SA = single addition). Data show growth of strains under SA conditions relative to M9G with all amino acids. All wildtype and mutant strains grew as expected based on their prototrophic/auxotrophic genetic background, thus validating our experimental approach. There was one exception: All *E. coli* mutants were unable to grow when isoleucine was omitted from the medium. This pattern strongly suggests that the uncovered isoleucine dependency was already present in the E. coli wildtype of the KEIO collection.

**A**

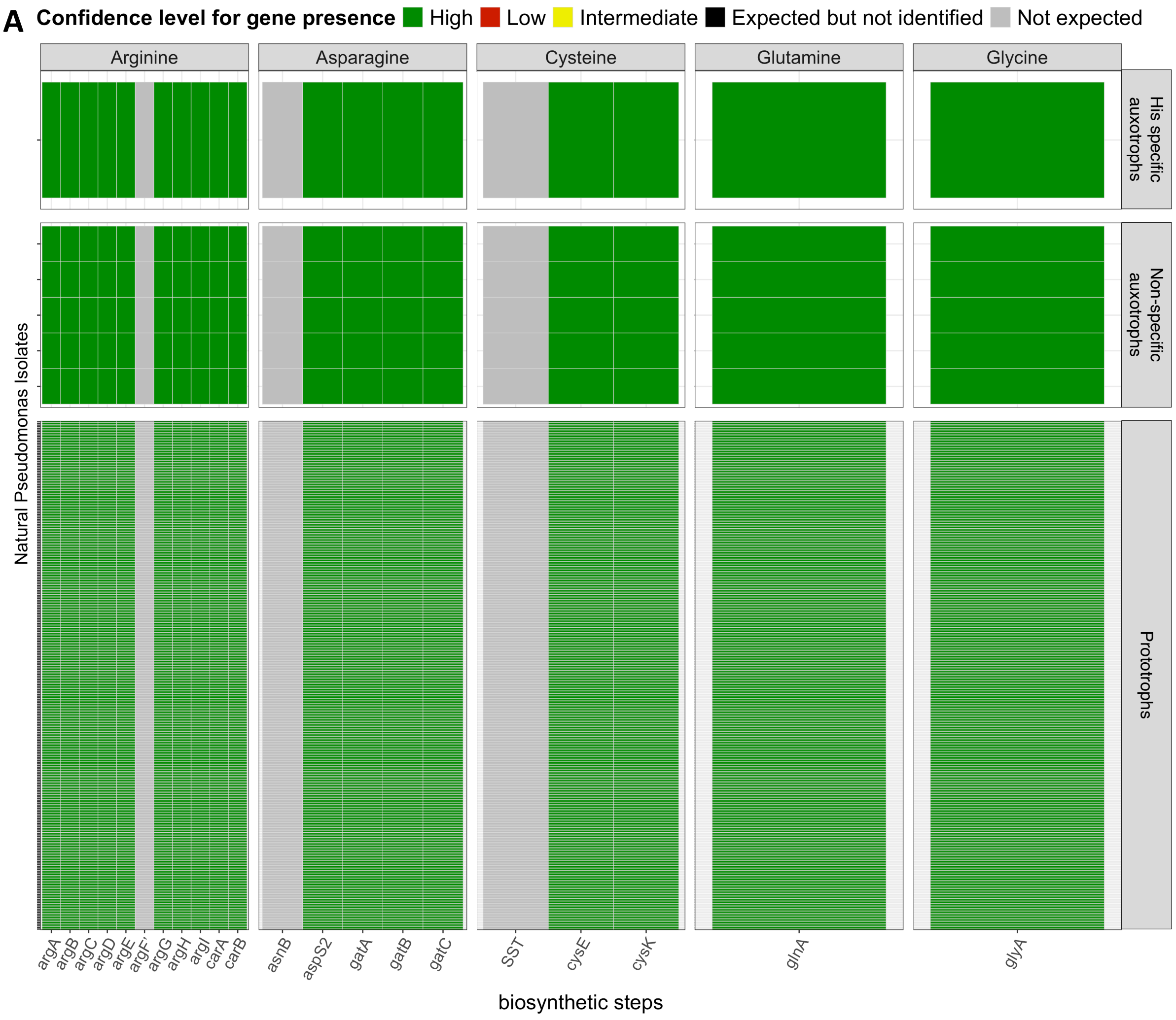
Confidence level for gene presence
High
Low
Intermediate
Expected but not identified
Not expected
Arginine
Asparagine
Cysteine
Glutamine
Glycine
His specific auxotrophs
Non-specific auxotrophs
Prototrophs
Natural Pseudomonas Isolates
argA
argB
argC
argD
argE
argF
argG
argH
argI
carA
carB
asnB
aspS2
gatA
gatB
gatC
SST
cysE
cysK
glnA
glyA
biosynthetic steps

B

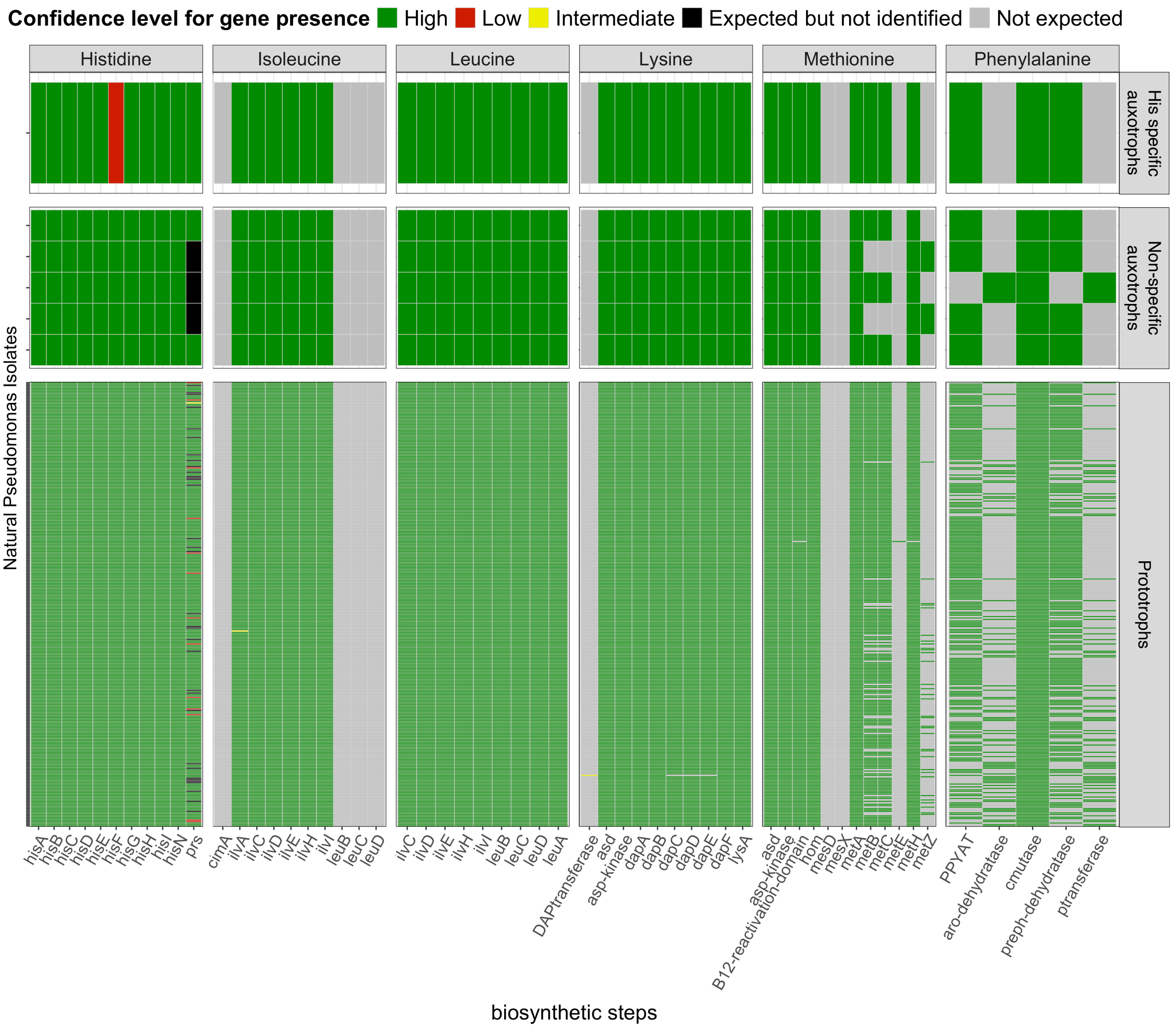
Confidence level for gene presence
High
Low
Intermediate
Expected but not identified
Not expected
Histidine
Isoleucine
Leucine
Lysine
Methionine
Phenylalanine
His specific auxotrophs
Non-specific auxotrophs
Prototrophs
Natural Pseudomonas Isolates
hisA hisB hisC hisD hisE hisF hisG hisH hisI hisN prs
cimA ilvA ilvC ilvD ilvE ilvH ilvI leuB leuC leuD
ilvC ilvD ilvE ilvH ilvI leuB leuC leuD leuA
DAPtransferase asd asp-kinase dapA dapB dapC dapD dapE dapF lysA
asd asp-kinase B12-reactivation-domain hom mesD mesX metA metB metC metE metH metZ
PPYAT aro-dehydratase cmutase preph-dehydratase ptransferase
biosynthetic steps

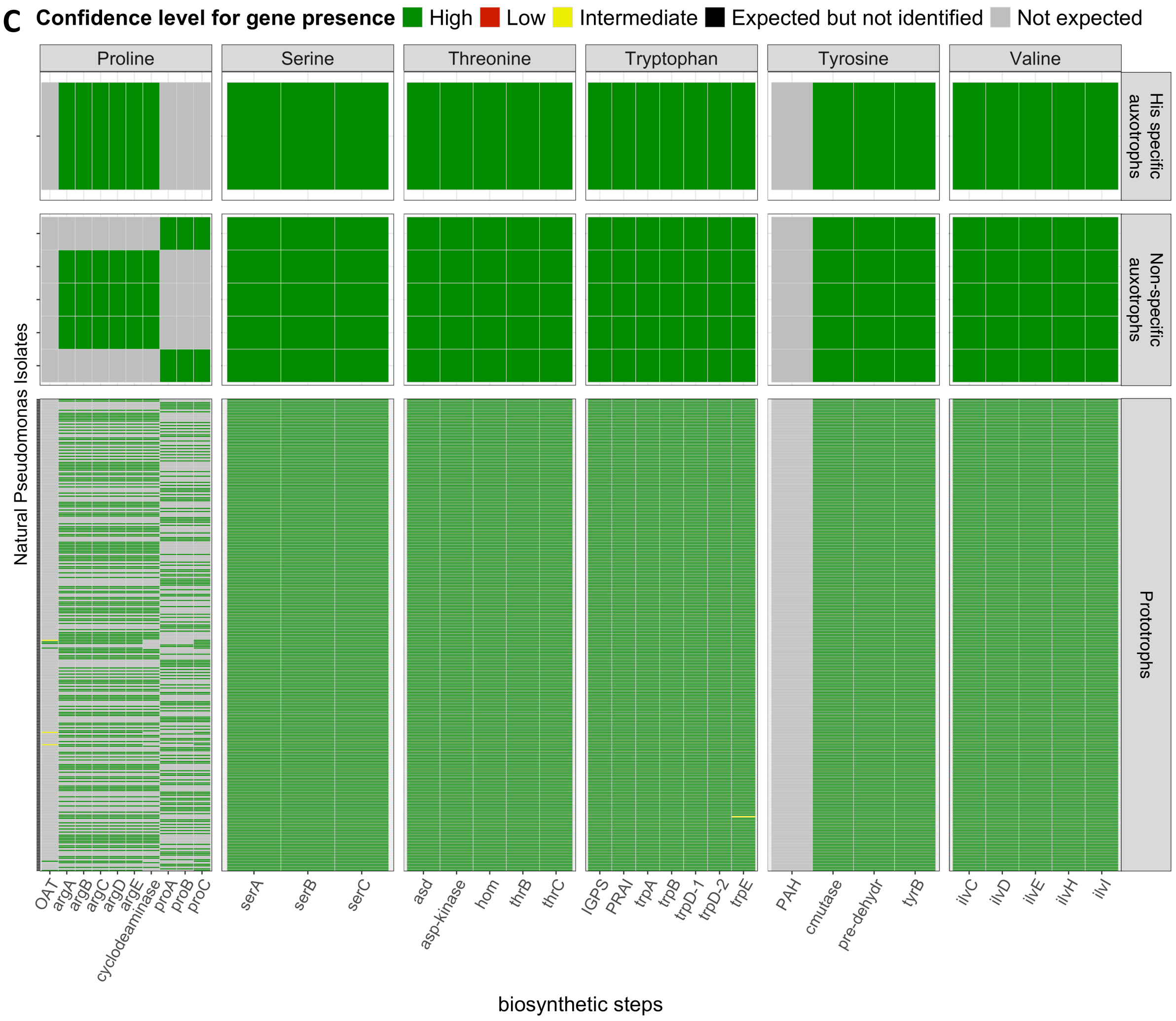


**Figure S2: Detailed GapMind predictions of amino acid biosynthesis pathway based on Prokka annotation.** Biosynthetic pathways for 17 amino acids predicted by GapMind for 314 natural *Pseudomonas* isolates: **A)** Arginine, Asparagine, Cysteine, Glutamine and Glycine; **B)** Histidine, Isoleucine, Leucine, Lysine, Methionine and Phenylalanine; **C)** Proline, Serine, Threonine Tryptophane, Tyrosine, Valine. Natural isolates (y axis) in each panel are separated into 3 categories based on experimental results (top: histidine specific auxotroph, middle: 5 non-specific auxotroph, bottom: prototroph). Each amino acid biosynthetic pathway is broken down into a series of genes (x-axis) forming a complete biosynthetic pathway. When a pathway is detected by Gapmind, it assigns a color representing the confidence level with which every gene involved in the detected pathway are found (green = high confidence, red = low confidence, yellow = intermediate confidence). When a pathway is detected but an essential gene is missing (below low confidence), it will be labeled as “Expected but not identified” (black). When a pathway is not detected, individual genes will not be searched by GapMind and will be labeled as “Not expected” (grey).

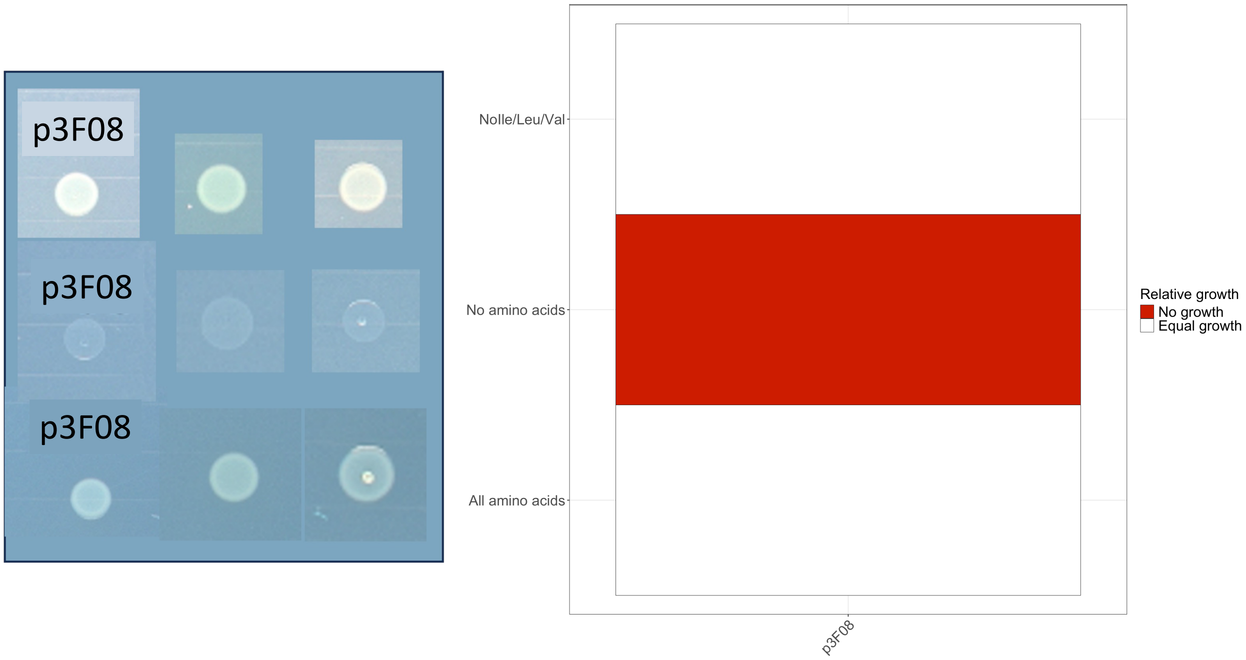


**Figure S3: Auxotrophy test for isolate p3F08 after GapMind predicted an auxotrophy.** GapMind predicted a missing gene (*ilvC*) central for the synthesis of isoleucine (Ile), leucine (Leu) and valine (Val) in isolate p3F08 with the PGAP annotation (see Fig. 3 in the main text). This suggest a complex case of auxotrophy that cannot be resolved with our SA/SO media approach. Thus, to test whether this specific GapMind prediction is correct or a false positive, we grew p3F08 in triplicates (left panel, columns) in M9G containing either all amino acids (bottom row), no amino acids (middle row) or a mix of 17 amino acids lacking isoleucine, leucine and valine (top row). We found that the growth of p3F08 in medium lacking the three amino acids was equal to its growth in the medium supplemented with all amino acid. We therefore concluded that GapMind prediction is false.

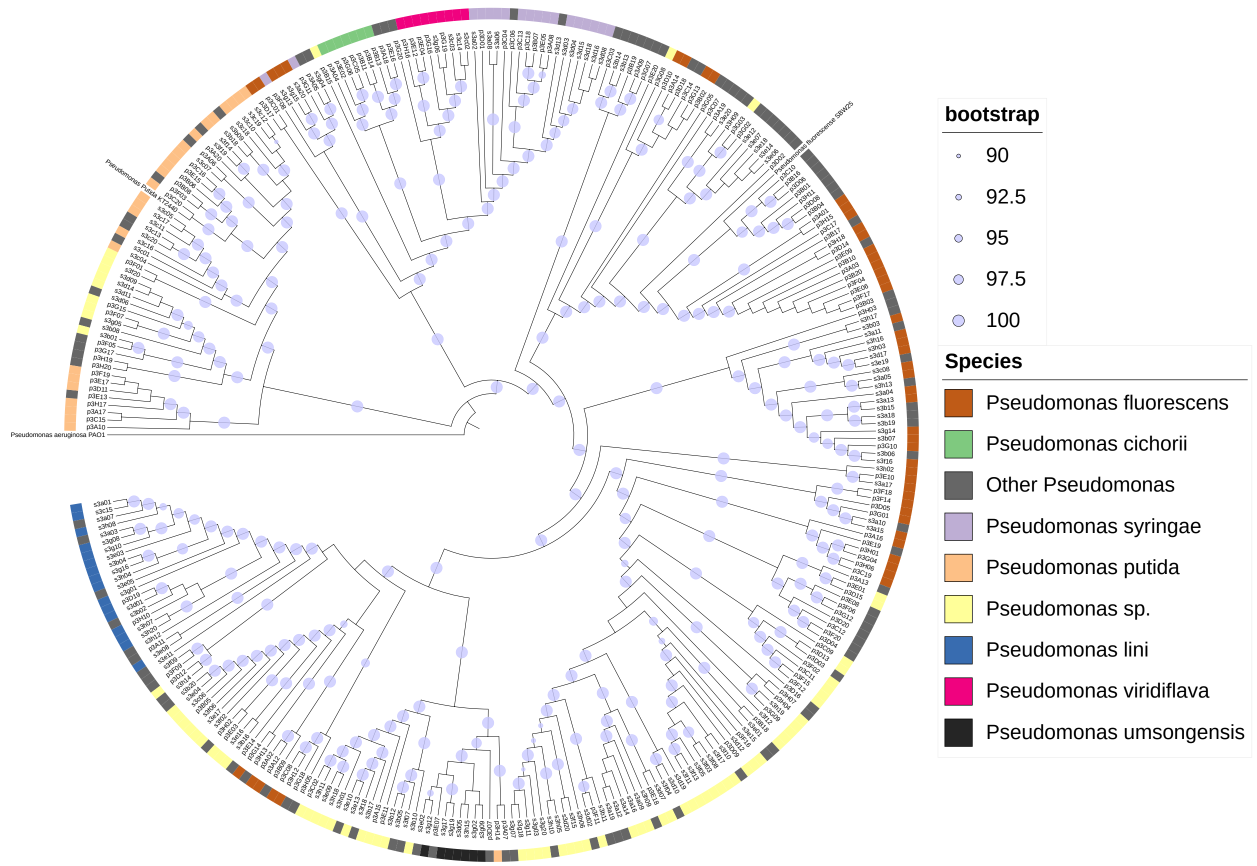


**Figure S4: Phylogenetic tree of the 314 natural *Pseudomonas* isolate with main represented species groups.** Phylogeny was constructed using 1240 single copy orthogroups (from Orthofinder) [1]. Allocation to species groups was determined using Kraken [2] using whole genome data and was carried out by MicrobesNG. Note that Kraken taxonomic allocation comes with substantial uncertainty. While the affiliation of isolates to the *Pseudomonas* genus occurred with high confidence (91% ± 6%), the species affiliation was associated with much lower confidence (52% ± 28%).
